# Global convergence of dominance and neglect in flying insect diversity

**DOI:** 10.1101/2022.08.02.502512

**Authors:** Amrita Srivathsan, Yuchen Ang, John M. Heraty, Wei Song Hwang, Wan F.A. Jusoh, Sujatha Narayanan Kutty, Jayanthi Puniamoorthy, Darren Yeo, Tomas Roslin, Rudolf Meier

## Abstract

Most of arthropod biodiversity is unknown to science. For this reason, it has been unclear whether insect communities around the world are dominated by the same or different taxa. This question can be answered through standardized sampling of biodiversity followed by estimation of species diversity and community composition with DNA sequences. This approach is here applied to flying insects sampled by 39 Malaise traps placed in five biogeographic regions, eight countries, and numerous habitats (>220,000 specimens belonging to >25,000 species in 463 families). Unexpectedly, we find that 20 insect families account for >50% of local species diversity regardless of continent, climatic region, and habitat type. These consistent differences in family-level dominance explain two-thirds of variation in community composition despite massive levels of species turnover, with most species (>97%) in the top 20 families encountered at a single site only. Alarmingly, the same families that dominate global insect diversity also suffer from extreme taxonomic neglect with little signs of increasing activities in recent years. Tackling the biodiversity of these “dark taxa” thereby emerges as an urgent priority because the arthropod groups comprising most of the global flying insect diversity are particularly poorly known.

## INTRODUCTION

Biodiversity loss is now widely recognized as a major threat to planetary health^1,2,3^. Halting the loss requires that the basic building blocks of biodiversity are known, so that changes can be recorded, external drivers identified, and appropriate policy actions be implemented. However, much of the terrestrial animal diversity belongs to hyperdiverse invertebrate clades that are so poorly known^4,5^ that it is difficult to obtain this critical information. For example, only 0.17G of the 2.16G records in the Global Biodiversity Information Facility pertain to arthropods while 67% relate to birds, although they only account for 10,000-20,000 species (0.2%) of the estimated 8-10 million multicellular species worldwide^6,7^. These numbers alone reveal the size of the knowledge gap for truly diverse taxa.

To allocate resources for discovering and conserving species, it is crucial to establish the relative contribution of different taxa to overall biodiversity so that the most diverse and abundant taxa can be given special attention. Identifying these taxa is furthermore important for understanding the structure of the living world and gaining insights into how community composition is shaped by evolutionary, biogeographic, or ecological factors^8^. Such analyses have been carried out for plants and snakes^9^ and revealed that, for example, a few snake clades dominate communities across the world^10^. Unfortunately, corresponding information is lacking for arthropods although they are found worldwide, functionally important^11^, and currently undergoing major changes in diversity and abundance^12,13^.

In this study, we analyze global taxonomic patterns among flying insects caught by Malaise traps. These standardized traps are widely used in global biomonitoring programmes, because they are efficient tools for collecting flying insects and semi-aquatic taxa^14–16^. However, Malaise traps are so effective that sample processing is a major challenge due to high specimen and species yields^14,17^. In addition, most specimens belong to species that cannot be identified because taxonomic expertise for hyperdiverse taxa is non-existent and/or dwindling for many species-rich clades caught in Malaise traps (“dark taxa”)^18^. Fortunately, recent advances in large-scale DNA barcoding with new sequencing technologies allow for processing large numbers of specimens rapidly and cost-effectively^19,20^. These data can then be converted into estimates of species diversity without formal description using molecular species delimitation methods and most species can be assigned to major insect clades for analysis of community structure.

We here determine the global taxonomic composition of Malaise trap samples^21^ from five biogeographic regions, eight countries, and very diverse habitats (>220,000 specimens belonging to >25,000 species living in habitats ranging from temperate meadows to tropical rainforests). We find surprising congruence with regard to which 20 insect families are dominant components of flying arthropod communities worldwide (>50% of species and specimens in samples). When we compare family-specific diversity with taxonomic attention and find that most of the particularly diverse and abundant taxa are poorly known and suffer from persistent taxonomic neglect. In other words, a very large proportion of the terrestrial animal biodiversity is not only unknown to science, but will also remain so for the foreseeable future unless such “dark taxa” become a preferred target for biodiversity science.

## RESULTS

Our study comprises 225,266 barcoded arthropods belonging to 463 families. They represent the insect diversity obtained from 39 traps across eight different sites and all continents excluding Australia and Antarctica. Applying a species-delimitation threshold of 3% sequence similarity^20,22^ reveals that the different Malaise traps yielded anywhere from 69 to 3426 molecular Operational Taxonomic Units (mOTUs) per trap. When these mOTUs were assigned to higher arthropod clades, we found that on average, 57.2% and 19.0% of the species in a trap belonged to the Orders Diptera and Hymenoptera (Supplementary Fig 1). When examined at family level, 61.7% of the specimens in a trap and 51.9% of the species in each trap belong to a set of only 10 insect families (henceforth referred to as the “top 10 families”). The next 10 families contributed an additional 9.7% of and 12.2% of specimens and species, respectively (Fig 1: see the “top 20 families”). Nearly one-fifth of the species per site (average 19.98%) belonged to only one dipteran family (Cecidomyiidae) (Fig 2A).

**Figure 1.**
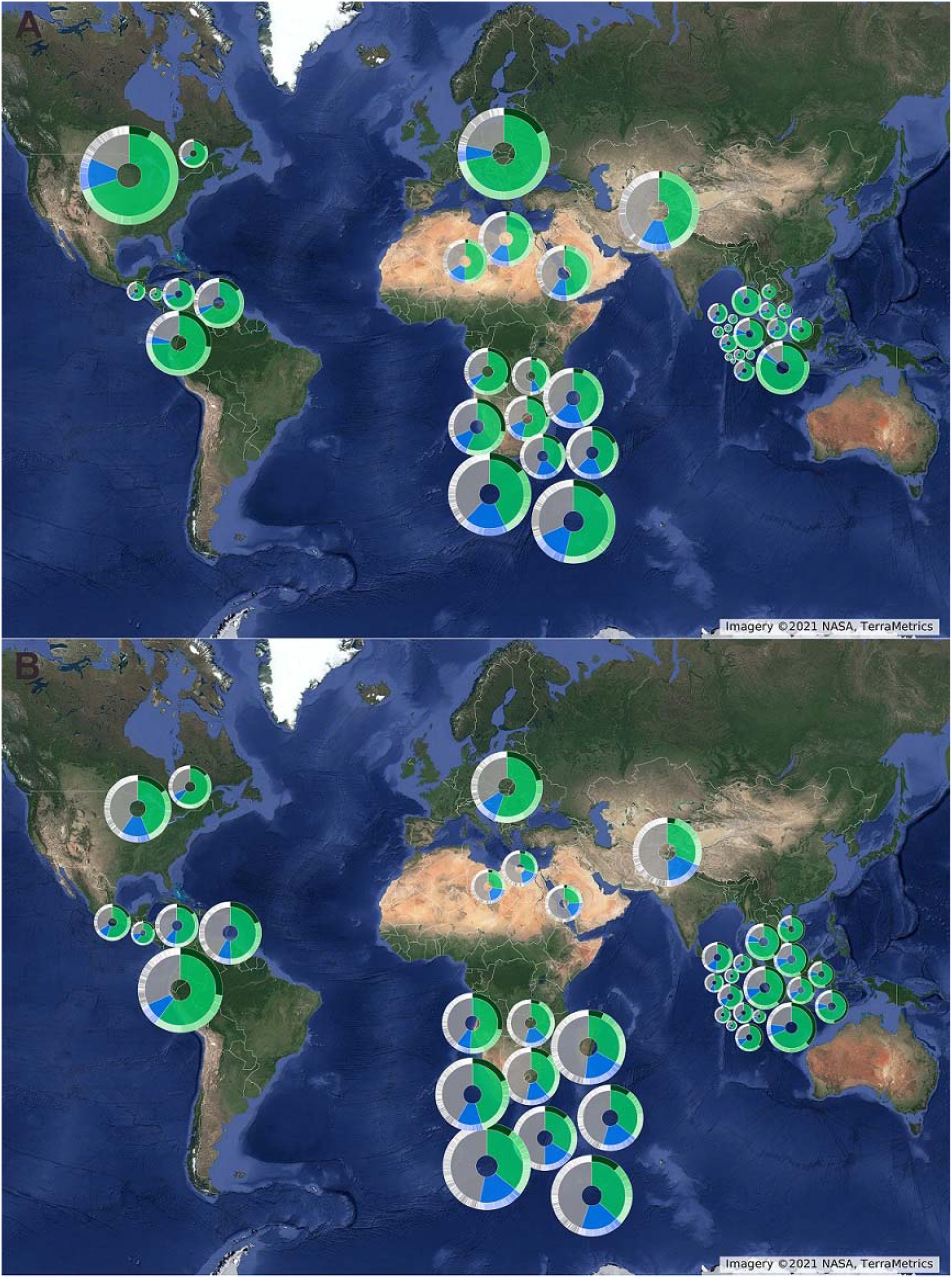
Global congruence in community composition of insects at the family-level (top: abundance; bottom: species). Each circle shows the taxonomic distribution of a sample obtained by one Malaise trap at a specific site. The top ten taxa are green, while the next ten are blue. The outer ring is divided by families, with the dark green margin representing the globally dominant family Cecidomyiidae (Diptera).

**Figure 2.**
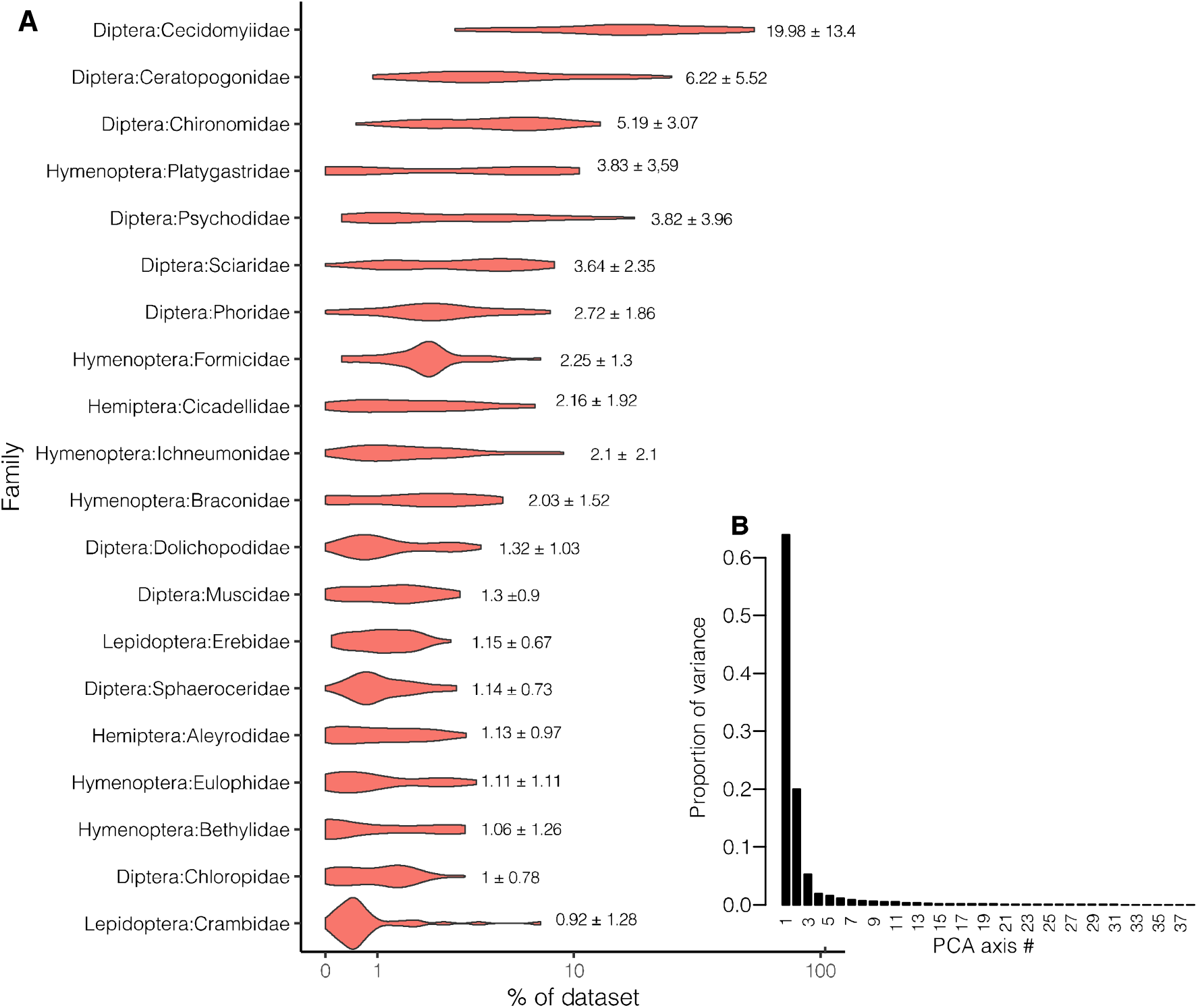
Global consistency in community composition of insects as shown by (A) proportional (%) species richness of the top 20 families among sites (x-axis transformed as log scale) and (B) proportion of variance absorbed by the first axis in a principal component analysis of variation in the proportion of species richness per insect family. Figures next to individual violins show the average proportional contribution ± SD for each individual family.

The relative species richness of individual insect families showed remarkably similar patterns across the globe, with the top 20 families (Fig 1, bottom panel) accounting for 41.2-72.3% of the total species richness regardless of continent or climate (Canada: 63.9%, Egypt: 47.3%, Germany: 65.8%, Honduras: 65.5%, South Africa: 54.5%, Pakistan: 49.4%, Saudi Arabia: 41.2% and Singapore: 72.3%). Yet, the high species richness of these families is not correlated with clade age (r=0.004; n=10023, p=0.69, Supplementary Fig. 5).

The evidence for this global pattern is perhaps best summarized by the consistent differences in species richness among families. In our main dataset (39 Malaise traps), 66.4% of the variation among traps in log-transformed species richness was explained by family. These results remained consistent irrespective of the species delimitation method used (objective clustering at a 3% distance threshold: 66.4%^23^ or ASAP^24^: 64.9%, based on adj. *R*^*2*^). In our expanded dataset (see Methods), which retained sample-level resolution for traps placed in Germany and Canada (56 datasets), the corresponding figure was 67.2% (objective clustering at 3%). The only qualitative difference in results between the main and extended data sets was whether Mycetophilidae (Diptera) and Crambidae (Lepidoptera) were included in the top 20. This list of top 20 families is furthermore robust to change when the next ten families are fused with their sister clades to account for disagreements on rank (Supplementary Table 3). Nineteen of the top 20 families remain in the list and the only change involved the replacement of one lepidopteran clade (Crambidae with Gelechiidae+Cosmopterigidae). Unsurprisingly, the global convergence of taxonomic composition is even stronger at order-level (90.6% of log-transformed species richness, Supplementary Figure 1).

The global convergence of relative species richness was also evident from a principal component analysis, which revealed that >60% of the variance can be explained by a single principal component (Fig 2B, Supplementary Figs 2B, 3B). In contrast, analysis of species turnover between the major regions shows that almost all species in the top 20 families (97.6%) are found in one site only (Supplementary Table 2). In other words, global variation in community composition is largely attributable to variation in the relative contribution of distinct species from a small set of families.

Given the disproportionate contribution of a few families to overall insect diversity across the world, we next examined whether the globally-dominant taxa attract appropriate global taxonomic attention. To characterize taxonomic attention or neglect, we first defined a “neglect index” (*NI*) as the ratio between the number of mOTUs found across the Malaise traps for a given family and the total number of species described as listed in the Catalog of Life [CoL: https://www.catalogueoflife.org/]. An NI value of 1 signals that we detected as many mOTUs in the only 39 traps as has been formally described for the entire world, whereas a low NI value will reveal that we found only a tiny proportion of all species described to date.

This analysis detected a positive correlation between the neglect index and the log number of mOTUs detected in our samples (main dataset: r=0.75, n=20, p= 0.00015; expanded dataset: r=0.76, n=20, p= 0.00011) (Fig 3, Supplementary Fig. 4) and a strong negative relationship between the neglect index and the number of species described per decade between 1980 and 2019 (main dataset, r=0.44, n=80, p= 3.2e-05, expanded dataset, r=0.48, n=80, p= 7.6e-06). In other words, the more neglected a taxon is, the fewer new species are described per decade. The slope of this relationship shows no improvement over time, as interaction decade×NI was far from significant (main dataset; anova: F=0.86, p= 0.53, expanded dataset, anova; F=0.80, p=0.58). Moreover, the number of taxonomists involved in monographic work (describing >=50 species in a decade) shows no increase over time (Fig 3c). In fact, the number of monographic revisions targeting the top 20 families was the lowest for decade of 2010-2019 (Fig. 3c: see the white part of the columns).

**Fig 3.**
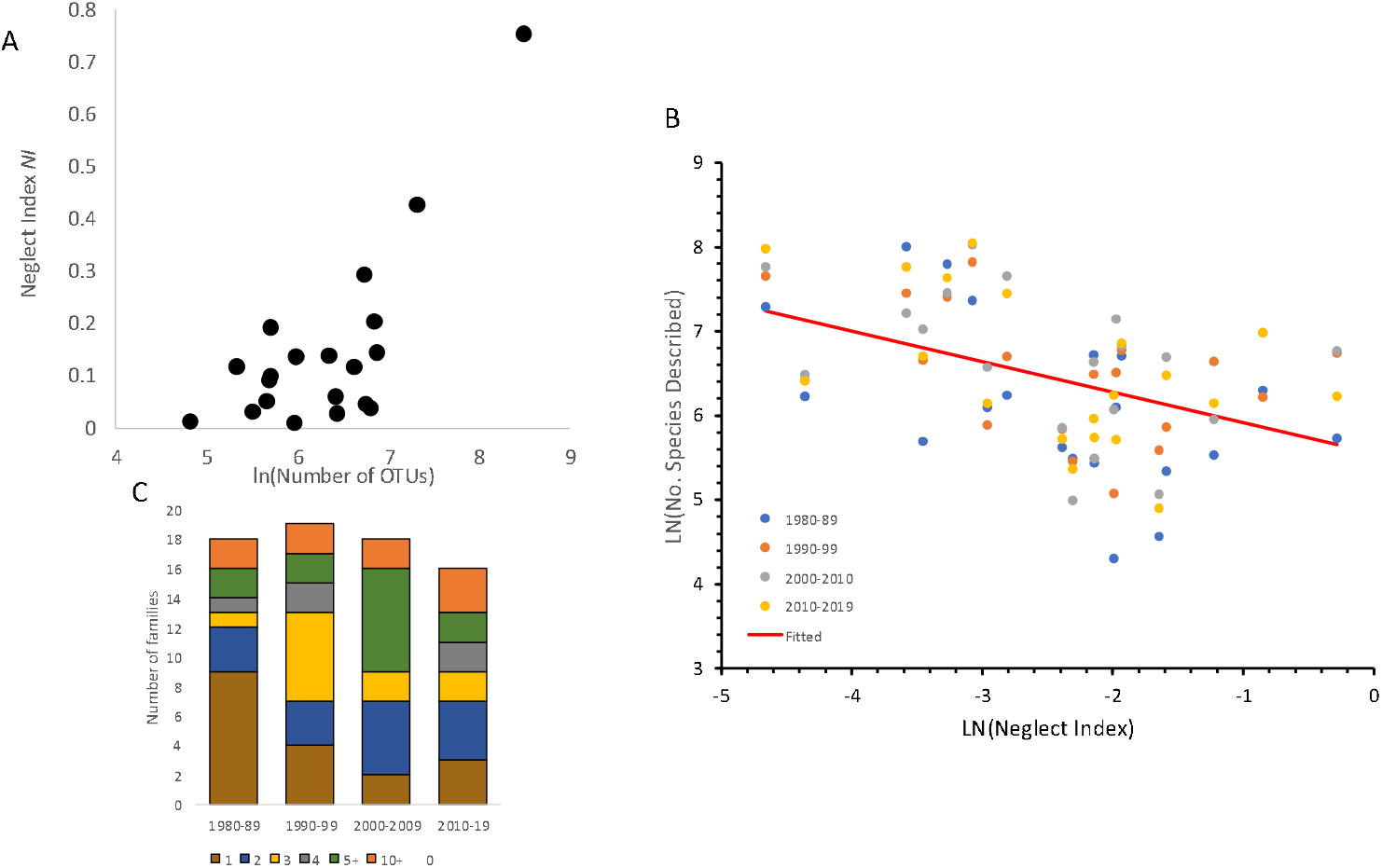
Taxonomic attention received by top 20 families shows no improvement over time. (A) The more species-rich a family is, the more it is neglected, and (B) the more neglected is a family, the less species have been described. (C) No indication of increasing taxonomic attention to the top-20 families over time. The bar chart shows the proportion of families (out of 20) that have been the focus of dedicated taxonomic work. As an indicator of taxonomists invested in a particular taxon, we calculated the number of individual authors describing at least 50 species per decade. The bar sections show the proportion of families with 0 (white),1 (brown), 2 (blue), 3 (yellow), 4 (grey), 5-9 (green) and 10+ (orange) such dedicated authors. (For the corresponding figure for the expanded dataset, see Supplementary Fig 4.)

## DISCUSSION

Arguably, insects remain the key taxonomic challenge for understanding animal diversity given that more than 80% are undescribed. We here reveal that the same 10-20 families dominate communities of flying insects around the world. This convergence is truly remarkable, given that the samples were collected across several distinct climatic zones and habitat types, including tropical rainforests, montane forests, cedar-savannah, bushwillow woodlands, thorn veld, mangroves and marshes. The biodiversity challenge posed by these “dark taxa” is formidable, given how species-rich they are at each site, and how high the species turnover is across sites. A good example are Cecidomyiidae (gall midges) which are globally hyperdiverse, dominate insect communities in terms of species counts and abundance *and* yet have received very little taxonomic attention.

We find that two thirds of the variance across Malaise trap samples is explained by family-membership of a species. This raises the question of how such a pattern can emerge across widely different ecosystems and large geographic scales. One explanation could have been that dominant insect clades ranked as families were older, and thus had more time for diversification. However, we detected no correlation between species richness per family and clade age (r=0.004; n=10023, p=0.69, Supplementary Fig. 4). Frequent dispersal by some families is also not a viable explanation because fewer than 3% of species in the top 20 families are found at multiple sites (1-9% in any given family, Supplementary Table 2). Instead, the convergence of taxonomic composition must be due to high diversification rates of taxa ranked above the species and below the family-level. For instance, the subfamily Cecidomyiinae contains 74% of the described species diversity of Cecidomyiidae and Psychodinae 59% of Psychodidae (CoL). Similarly, 45% of described diversity of ants (Formicidae) is in the subfamily Myrmicinae. It will be important to identify whether the same subclades in the hyperdiverse families are responsible for the high species diversity across biogeographic regions and habitats. Such analysis has been carried out for snakes, where a few rapidly diversifying clades dominate globally^10^. These clades (e.g., Colubrinae) are apparently better colonizers and are successful in geographically disparate regions regardless of the presence of other taxa. However, compared to most insect clades in our study (mean= 158 mya), colubrines are very young (<50 mya) and the insect patterns reported here evolved over a much greater span of time. The next step in understanding this pattern is the detailed analyses of diversification rates and colonization events for clades below the family and above the species level for the globally dominant taxa.

Our study analyzes insect communities captured in Malaise traps. These traps are particularly effective for sampling flying insects ^16,25^ and used widely in many large scale insect biomonitoring programmes^14,15^ because they yield standardized samples that are useful for assessing site-to-site differences. However, Malaise trap samples will contain few canopy species or taxa that are best sampled with other techniques such as flight intercept traps or targeted sweep-netting^26 27^. It will thus be important to repeat similar comparative and global analyses of insect communities based on samples obtained with alternative sampling techniques. This would reveal to what extent the order-level dominance of Diptera and Hymenoptera^28^ is so widespread that they are indeed more species-rich than Coleoptera and Lepidoptera^29^. Note that among the twenty most common families identified in the current study, ten are dipteran (Cecidomyiidae, Ceratopogonidae, Chironomidae, Psychodidae, Sciaridae, Phoridae, Dolichopodidae, Muscidae, Sphaeroceridae and Chloropidae) and six are hymenopteran (Platygastridae, Formicidae, Braconidae, Ichneumonidae, Eulophidae and Bethylidae) (Fig 2A).

Taxonomic impediments are a major reason why many biodiversity community patterns have yet to be resolved. What allowed us to here address the global community structure of flying insects at species and family level was a combination of new high-throughput species discovery methods via large-scale barcoding^30,31^ and efficient taxonomic assignment techniques. These new tools will be essential for addressing taxonomic biases that have interfered with studying, understanding, and explaining biodiversity patterns across all of animal diversity. As biodiversity loss is threatening environmental health globally, obtaining unbiased biodiversity information across all taxa is crucial. Such unbiased information will complement the large amount of data on large and charismatic species^32^ and lead to biodiversity assessments that include those taxa that dominate ecosystems in terms of species diversity, abundance and biomass^33^.

Arguably, one of the most worrying findings of our study is the taxonomic neglect of the most important insect families. We found that the more a family contributed to insect communities around the world, the more it has been neglected. Furthermore, we found no evidence for any increase in the taxonomic attention over time. On the contrary, the neglect proved most severe for the most diverse insect families and there is no evidence that they have been receiving more attention in the last decade. This seems a major shortcoming of modern biodiversity science.

The neglect of dominant taxa also compromises current estimates of global species richness. These types of estimates frequently use ratios of species richness across families (e.g., ratio of butterflies diversity to known insect diversity in Britain^34^). Such estimates are severely affected if the richness estimates for dark taxa are incorrect. For instance, in UK, only 2.7% of described insect diversity belong to Cecidomyiidae^35^, compared to an average of 20% found in the Malaise trap communities analysed here. Assuming that the true diversity of Cecidomyiidae is closer to 20%, the global species richness estimate would shift from 5.4-7.2 million to 6.5-8.7 million species although we are here only revising the diversity estimate for one dark taxon (see Supplement Material 1). Thus, the neglect of dark taxa is very likely to also severely affect our perception of how life on Earth is organized. Hence, there is a need to start intensive work on dark taxa (“dark taxon biology”) to reveal Earth’s true diversity.

Overall, our study suggests that biodiversity research on 10 insect families which were discovered with comparatively new methods should be a global priority, but similar scalable and new approaches are also needed for species interaction research in order to understand the functional significance of the key taxa. This can be achieved through the implementation of a reverse-workflow^19^ to taxonomy, where all specimens are initially sorted to putative species based on barcodes.

Taxonomists can then revise species boundaries and describe species while molecular ecologists use the same vouchers for gaining insights into the biology of the species. Close collaboration can constitute a step change in the rate of species discovery in key clades, but this will be *conditional* on the adequate resources directed to priority taxa.

## MATERIALS AND METHODS

### Datasets, sample collection and processing

The study used DNA barcode generated for full Malaise trap samples across the globe. New datasets were generated for 18 samples from different habitat types in Singapore. These covered a terrestrial forest (5 traps), a mangrove forest (7 traps), coastal forests (3 traps), and a marsh (3 traps). The samples were recovered weekly between 3 May 2019 to 9 May 2019. Specimens were preserved in molecular-grade ethanol and then barcoded. The remaining data came from published studies that sequenced all specimens from a Malaise trap samples. This included datasets from Germany^15^, Canada ^36,37^, South Africa ^38^, Pakistan^39^, Saudi Arabia^39^, Egypt^39^ and Honduras^40^ (Supplementary Table 1). In order to avoid strong geographic biases in our study, we used only data for 9 of the 20 Malaise traps from South Africa (Kruger National Park), each representing a different habitat. For the study by Telfer et al. (2015)^37^ for Malaise trap sequencing in Canada, we limited analysis to the largest sample.

### DNA sequencing, barcoding and identification

Insects from Malaise traps placed in Singapore were processed in a similar approach to Yeo et al. (2021)^20^ and Srivathsan et al. (2021)^30^ in that we used NGS (Next Generation Sequencing) barcoding methods^41^. Briefly, DNA was extracted using 10-30 μL hotSHOT ^42^ per specimen and heated to 65°C for 18 min, followed by 98°C for 2 minutes, after which an equal volume of neutralization buffer was added. A 313-bp fragment of *cox1* was amplified using primers mlCO1intF and 5′- GGWACWGGWTGAACWGTWTAYCCYCC-3′^43^ and jgHCO2198: 5′- TANACYTCNGGRTGNCCRAARAAYCA-3′^44^. The primers were tagged with a 13-bp tag at the 5’ end as described by Srivathsan et al.^30,17^ PCRs were conducted in 96*-* well plates using one negative control per plate, and each PCR mix contained 7□μl Mastermix (CWBio), 0.5□μl BSA (1□mg/ml), 0.5 μl MgCl_2_ (25 mM), 1□μl each of primer (10□μM) and 4□to 7 μl of template DNA. The cycling conditions were: 5□min initial denaturation at 94□°C followed by 35□cycles of denaturation at 94□°C (30□s), annealing at 45□°C (1□min), extension at 72□°C (1□min), and followed by final extension of 72□°C (5□min). A subset of 8-15 products per plate were run in agarose gels to assess PCR success. Samples were pooled and purified using Ampure XP beads (Beckman Coulter). Pooled samples were sequenced either using Illumina HiSeq 2500 (2×250-bp) or using MinION sequencer (Oxford Nanopore Technologies). Illumina sequencing was outsourced while MinION sequencing was conducted in-house using an R9.4 flowcell. Libraries were prepared using the SQK-LSK109 Ligation Sequencing Kit based on the modified protocol described^30^. Fast basecalling model as implemented in Guppy was used as high accuracy basecalling was not available at the time of data-processing.

Data analysis was conducted based on procedures described by Wang et al. (2018)^19^ for Illumina reads. Here, paired-end reads were merged using PEAR^45^. Reads were demultiplexed allowing for up to a 2 bp mismatch in primer sequences while no mismatch was allowed in the tag sequence. Demultiplexed reads for each specimen were merged to form unique sequences, and only datasets having at least 50 sequences were processed further. A dominant sequence was identified and if it had a read count exceeding 10, it was “called” as the DNA barcode for the specimen, as long as it was also at least 5 times as common as the second dominant sequence. MinION sequence data was processed using *minibarcoder*^46,17^ which both demultiplexes the data and calls the barcodes. The final consolidated barcode sets were used for further analysis.

Barcodes were clustered at 1% using objective clustering (see below) and specimens were sorted physically based on their cluster assignments. For Singapore samples, each cluster was morphologically identified to family. For Lepidoptera specimens, as well as for a small number of other specimens where morphological identification was not possible, we assigned specimens to families based on DNA characters alone. This was done by conducting BLAST against NCBI-*nt* database as well as searches against the BOLD Systems Identification engine (https://boldsystems.org/index.php/IDS_OpenIdEngine). A taxonomic assignment to family was accepted if there were no conflicting family-level matches in the top 20 unique matches. For all other morphologically identified specimens, DNA based identification was examined and any conflict with morphology was resolved through re-examination of morphology. If a conflict persisted, the specimen was not identified to family. Taxonomic classifications for published studies were based on metadata provided by the studies. These studies employed various methods of identifications including morphology, matches on BOLD, and tree-based identifications. It was noted that several Hymenoptera identified based on morphology as Scelionidae matched Platygastridae on BOLD Systems. This is likely due to recent classification changes. The “old” Platygastridae (prior to 2007) was treated as Platygastroidea until very recently when it has been split into several families^47^. We here follow several other recent studies^38,40^ that used the old circumscription of Platygastridae.

### Species delimitation

Prior to species delimitation, we excluded sequences that contained a stop codon when translated using the invertebrate mitochondrial genetic code, or which could not be identified to family. Secondly, in order to ensure that the large datasets had sufficient sequence overlap for multiple sequence alignments, short barcodes were excluded. Any sample/trap that contained <100 barcode sequences was excluded. For datasets containing 313-bp barcode sequences, the length cutoff was 300 bp, while that for datasets containing 658-bp barcodes was 500 bp. Barcodes were aligned using MAFFT v7^48^. Species delimitation was conducted using objective clustering, which is a distance-based clustering algorithm originally described in Meier et al. (2006)^49^. Species delimitations were also conducted using another distance based approach (ASAP)^24^ and a tree-based approach (Poisson Tree Processes or PTP)^50^. Most analyses initially used molecular Operational Taxonomic Units (mOTUs) obtained with objective clustering using a 3% distance threshold before testing the results with ASAP and PTP. We find that the results are very similar irrespective of clustering method or distance threshold (see Supplementary Table 1). Species delimitations were conducted for individual datasets independently. An estimate for total species diversity was obtained using USEARCH (v 11.0.667) ^51^ cluster_fast (-sort length -id 0.97) for the 225,266 sequences used in the study.

### Statistical analyses of community composition

All analyses were conducted in R v4.1.2^52^. Analyses were limited to insects (i.e., spiders, Collembola etc were excluded: see list in Supplementary Table 14 and 15). Overall 225,266 sequences were analysed. Two different datasets were analyzed: one where all sequences available for the same Malaise trap were combined (39 traps, main dataset) and one where the sample-level resolution for the datasets from Germany and Canada was retained (56 datasets, expanded dataset). This was feasible due to availability of high-quality metadata such that weekly samples could be treated separately. Community composition at the family-level was analysed using a linear model. Here, we first logarithmically-transformed the proportion of mOTUs for each dataset, adding 0.01 to zero proportions (since (log[0] is undefined).

We then modelled the transformed response variable as a function of family [*lm(log(Proportion+0*.*01)∼Taxon,data=dataset)*], using adjusted *R*^*2*^ values as the key statistic of variance explained. Furthermore, we ran a PCA (Principal Component Analysis) on the community matrix of each site (with cell values equaling the proportion of species richness per family) using *rda*. We set scale as FALSE, and used a barplot of relative eigenvalues to assess the percentage variation explained by each principal component.

The top 20 families were identified based on ranking of average proportion of mOTUs per family. To test whether the subjective nature of family-ranks influence the results, we also examined which taxa are in the top 21-30 taxa. We then merged each with its sister clade based on recent phylogenies to test whether the merge would generate a taxon that would be included in the list of top 20 families. Next, we examined whether the high number of species in these clades across Malaise trap samples was due to high species-level dispersal rates. We analyzed species turnover across “sites”. “Sites” were broadly defined as Canada, Egypt, Germany, Honduras, Saudi Arabia, Pakistan, South Africa and Singapore.

Lastly, we assessed the correlation between the age of family and the proportion of mOTUs. Family ages were obtained from TimeTree (http://timetree.org, version 5). Ages for families lacking information in TimeTree were obtained from major large-scale studies involving dating^53–57^. For the various statistical analyses we excluded families with <10 specimens across all the samples and families that are present in one sample only.

### Assessment of taxonomic neglect

In order to assess how much taxonomic attention has been given to the families dominating Malaise trap samples, the total number of species described for each of the top 20 families was obtained from Catalog of Life v22.3 [CoL: https://www.catalogueoflife.org/]. Bethylidae, CoL lacked information although the family was listed. We thus used values from a recent checklist instead^58^. For two families, we used the species richness in superfamilies (Noctuoidea, Platygastroidea). This was either because the family was not listed (Erebidae) or because of recent changes in family-level classification (Platygastridae, see above). In order to understand whether a particular taxon is neglected, we defined a neglect index (*NI*) as *N*_*mOTU*_ /*N*_*sp*_, where *N*_*mOTU*_ is the number of mOTUs for a given family across the whole dataset and *N*_*sp*_ is the total number of species described as obtained from CoL. To test for a relation between diversity and taxonomic neglect, we then calculated the linear correlation between *NI* and ln(*N*_*mOTU*_). For each of the top 18 families and 2 superfamilies, species delimitation was conducted independently using objective clustering.

We next examined taxonomic attention to the top 20 taxa over time. We counted the species descriptions in *Zoological Record*. All data for the top 20 families were downloaded by search term ST=[Taxon name], as encompassing 181,985 studies. Studies in the last 4 decades (1980-2019) which describe species were identified by the s*p nov* epithet in the organism field. This identified 16,362 studies. The information on organism was then extracted to obtain the species and the family name, along with information on the year of publication and the authors involved. The data were then parsed to obtain the total number of species described in the study. For Erebidae, Crambidae, Aleyrodidae and Platygastridae, we assessed information at the level of the superfamily.

To test for a change in the relation between species diversity and neglect, we tested for an interaction between NI and Decade. To this aim, we compared two ANCOVA models fitted to the univariate data: lm(log(N_sp10_)∼ *NI*, data=UnivariateData), and lm(formula = Value ∼ *NI* * Decade, data = UnivariateData). Here, N_sp10_ is the number of species described in a decade, with decades being 1980-1989, 1990-1999, 2000-2009 and 2010-2019. The fit of the two respective models was then compared to each other by ANOVA (anova (model1,model2)).

Lastly, to understand whether taxonomic work dedicated to the top 20 families has changed over time, we extracted the information on the number of authors highly dedicated to a particular family, as scored from the number of authors exceeding a particular threshold (*S*_*threshold*_) of species descriptions. Given that some of these descriptions involved multiple authors, the score for each author *i* was calculated as 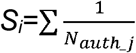, where *N*_*auth*__*j* is the number of authors in the study *j*. For an article in which two authors described a species, each author would thus get an author score of 0.5 for this species. The number of highly dedicated authors was then scored as the number of authors with S_*i*_> *S*_*threshold*_ per decade, where *S*_*threshold*_=50.

## ACKNOWLEDGEMENTS

Special thanks go to the team from the National Biodiversity Centre of NParks and the Mandai Park Holding for assistance and permits (Permits: NP/RP12-022-4, NP/RP12-022-5, NP/RP12-022-6). Financial support was provided by Mandai Park Holding, Ministry of Education Singapore, and Center for Integrative Biodiversity Discovery, Leibniz Institute for Evolution and Biodiversity Science, Museum für Naturkunde Berlin.

