## Supplementary Figures for "Global convergence of dominance and neglect in flying insect diversity"

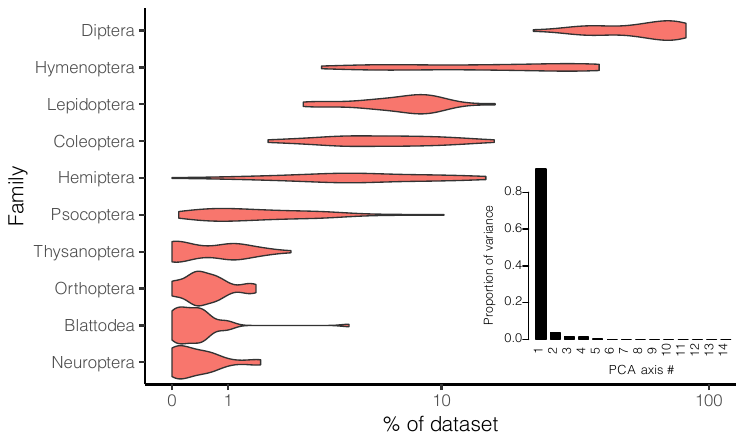

Supplementary Fig 1. (A) Proportion of species richness in top 10 Orders and (B) PCA analysis based on main dataset

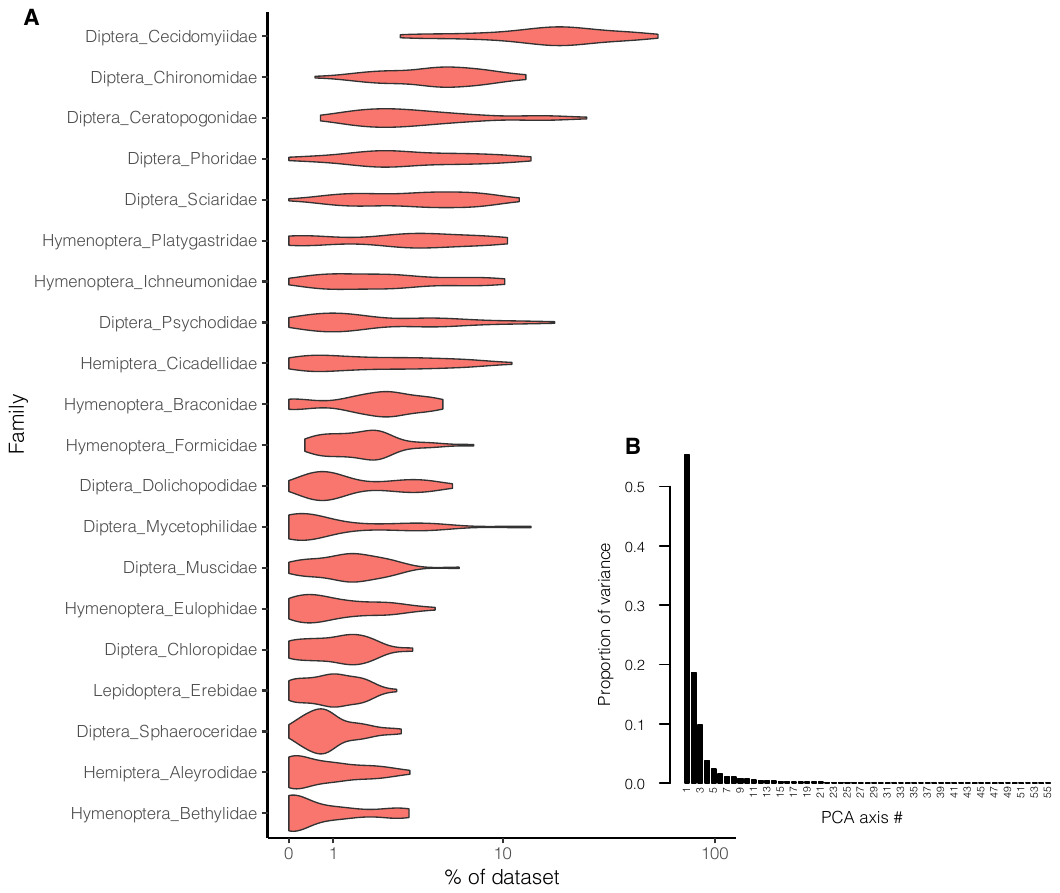

Supplementary Fig 2. (A) Proportion of species richness in top 20 families and (B) PCA analysis based on expanded dataset (See detailed legend of Fig 1)

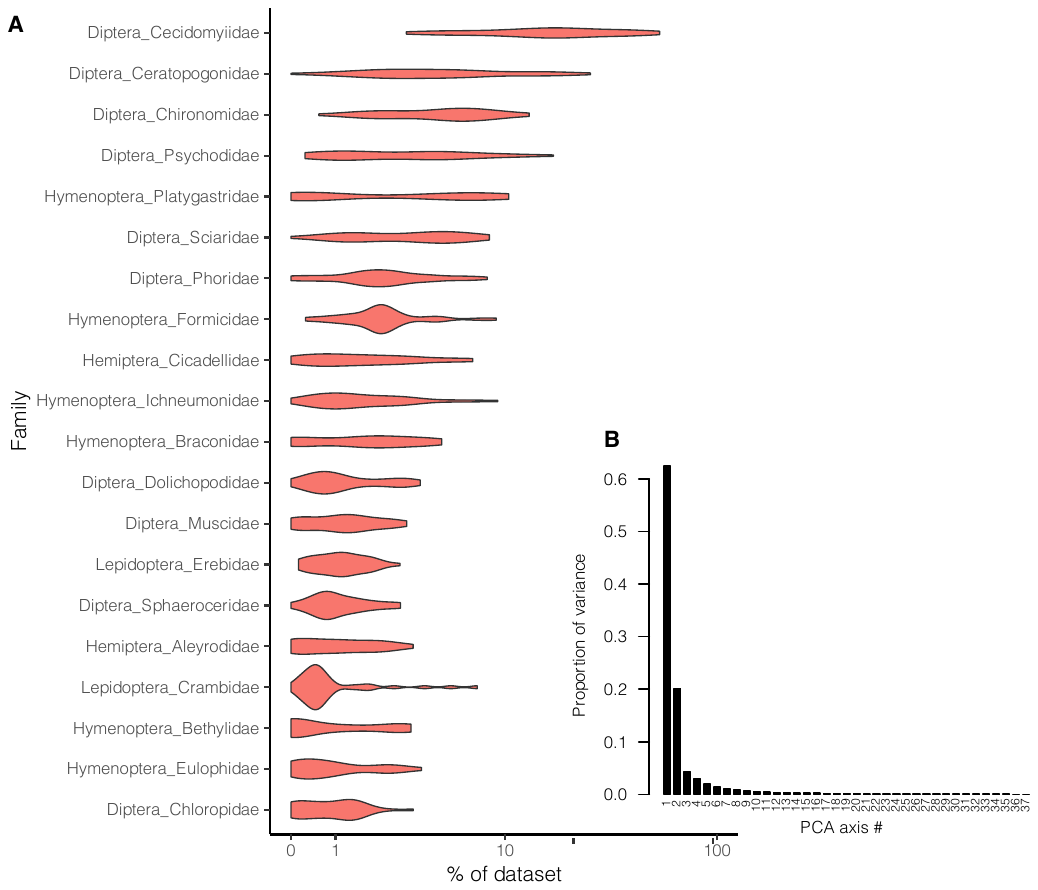

Supplementary Fig 3. (A) Proportion of species richness in top 20 families and (B) PCA analysis based on main dataset and ASAP species delimitation (See detailed legend of Fig 1). Algorithm failed for species delimitation for TrapW from dataset from South Africa, and this trap was excluded

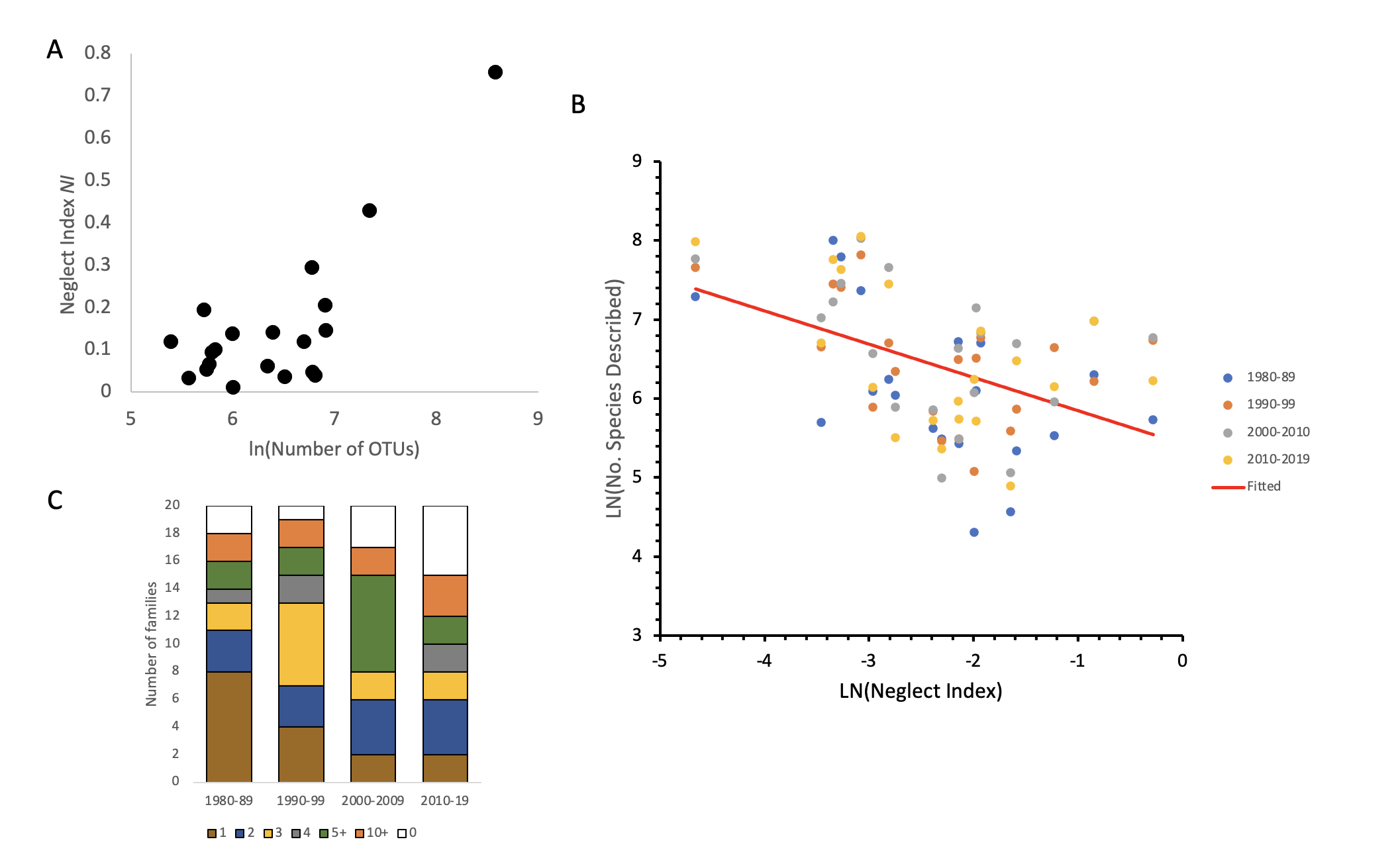

Supplementary Fig 4. Taxonomic attention received by top 20 families based on expanded dataset. (A) positive correlation between neglect index and log number of OTUs in the study (B) relationships between neglect index and number of species described in each decade (C)number of authors describing equivalent of 50 species per decade for each of top 20 families.

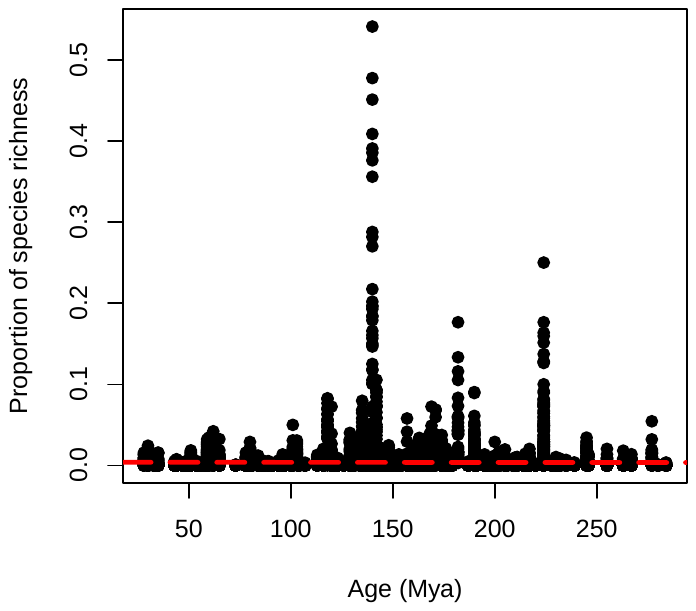

Supplementary Fig 5. Correlation between age of clade (Family) and proportion of species richness in the trap

Table 1: Number of mOTUs based on different species delimitation methods and distance thresholds

| **Trap** | **#specimens** | **#specimens after removing non flying insects** | **OC2%** | **OC3%** | **OC4%** | **ASAP1** | **ASAP2** | **ASAP3** | **MPTP** |
| --- | --- | --- | --- | --- | --- | --- | --- | --- | --- |
| **Singapore** | | | | | | | | | |
| CON03 | 147 | 145 | 79 | 79 | 78 | 78 | 78 | 83 | 80 |
| CON05 | 353 | 350 | 116 | 114 | 112 | 112 | 99 | 114 | 119 |
| KM03 | 1544 | 1432 | 432 | 425 | 420 | 411 | 417 | 409 | 431 |
| KM04 | 8533 | 8194 | 1282 | 1247 | 1227 | 1180 | 1112 | 1211 | 1271 |
| KM05 | 1623 | 1455 | 419 | 409 | 404 | 411 | 412 | 407 | 413 |
| MISL02 | 1150 | 1061 | 636 | 628 | 622 | 620 | 623 | 615 | 631 |
| MISL03 | 3156 | 2708 | 706 | 700 | 695 | 680 | 687 | 668 | 706 |
| MISL06 | 988 | 816 | 419 | 408 | 403 | 403 | 396 | 402 | 407 |
| MISL07 | 2386 | 750 | 396 | 390 | 386 | 376 | 2 | 410 | 402 |
| MISL10 | 2075 | 1879 | 574 | 560 | 550 | 551 | 535 | 557 | 566 |
| PU01 | 1418 | 1317 | 528 | 518 | 516 | 509 | 508 | 502 | 473 |
| PU22 | 383 | 359 | 205 | 201 | 198 | 197 | 182 | 200 | 225 |
| PU23 | 288 | 273 | 133 | 132 | 130 | 130 | 132 | 129 | 133 |
| PU24 | 545 | 493 | 222 | 218 | 215 | 214 | 213 | 209 | 221 |
| PU25 | 1193 | 1127 | 360 | 359 | 355 | 350 | 360 | 354 | 360 |
| PU27 | 554 | 493 | 250 | 247 | 245 | 245 | 247 | 243 | 249 |
| PU29 | 369 | 268 | 171 | 170 | 169 | 173 | 169 | 172 | 173 |
| PU26 | 4271 | 3193 | 984 | 971 | 959 | 903 | 903 | 925 | 983 |
| **Canada** | | | | | | | | | |
| 12-Sep-12 | 1572 | 1515 | 483 | 469 | 459 | 441 | 446 | 445 | 503 |
| 15-Aug-12 | 1131 | 1022 | 374 | 368 | 355 | 329 | 338 | 339 | 375 |
| 18-Jul-12 | 1273 | 1195 | 380 | 372 | 363 | 359 | 360 | 380 | 390 |
| 1-Aug-12 | 814 | 774 | 251 | 246 | 240 | 239 | 223 | 241 | 249 |
| 20-Jun-12 | 2440 | 2345 | 635 | 619 | 606 | 587 | 593 | 584 | 648 |
| 23-May-12 | 2275 | 2216 | 451 | 442 | 431 | 434 | 430 | 423 | 469 |
| 29-Aug-12 | 1685 | 1585 | 419 | 410 | 397 | 380 | 378 | 417 | 427 |
| 4-Jul-12 | 2111 | 2011 | 611 | 589 | 569 | 542 | 542 | 606 | 611 |
| 6-Jun-12 | 1931 | 1862 | 496 | 485 | 472 | 477 | 461 | 478 | 505 |
| 9-May-12 | 3795 | 3710 | 453 | 438 | 418 | 393 | 388 | 376 | 457 |
| DeWaard et al (all) | 27935* | 27143* | 2211 | 2121 | 2012 | 1888 | 1894 | 1894 | NA |
| Canada Site3 (Telfer et al.) | 2673 | 2550 | 925 | 901 | 880 | 883 | 847 | 886 | 941 |
| **Germany** | | | | | | | | | |
| 20-Jun-12 | 3194 | 3070 | 793 | 773 | 749 | 763 | 755 | 752 | 810 |
| 22-Aug-12 | 4529 | 4446 | 815 | 786 | 760 | 777 | 763 | 764 | 825 |
| 22-May-12 | 1691 | 1617 | 347 | 339 | 333 | 335 | 338 | 335 | 349 |
| 22-Sep-12 | 720 | 704 | 253 | 249 | 244 | 246 | 243 | 242 | 252 |
| 25-Jul-12 | 5094 | 5034 | 864 | 835 | 809 | 818 | 815 | 822 | 882 |
| 3-Sep-12 | 2366 | 2329 | 452 | 442 | 429 | 439 | 423 | 459 | 462 |
| 4-Jul-12 | 2234 | 2215 | 636 | 618 | 607 | 633 | 626 | 620 | 646 |
| 8-Jun-12 | 2531 | 2460 | 549 | 533 | 523 | 526 | 522 | 528 | 555 |
| 8-May-12 | 1646 | 1624 | 242 | 239 | 233 | 238 | 235 | 232 | 242 |
| Geiger et al. (All GMTPE) | 24005 | 23499 | 2502 | 2397 | 2284 | 2316 | 2337 | 2336 | NA |
| **South Africa** | | | | | | | | | |
| TrapC | 7437 | 6929 | 2048 | 1947 | 1862 | 1846 | 3025 | 1793 | 2034 |
| TrapE | 8546 | 7978 | 2262 | 2190 | 2125 | 2207 | 2103 | 2102 | 2279 |
| TrapI | 6662 | 6058 | 2148 | 2077 | 2016 | 2009 | 2009 | 2022 | 2180 |
| TrapJ | 11576 | 10466 | 2987 | 2885 | 2817 | 2815 | 2804 | 2822 | 3006 |
| TrapO | 6768 | 6045 | 1859 | 1791 | 1738 | 1588 | 2843 | 2452 | 1858 |
| TrapP | 4923 | 4291 | 1229 | 1197 | 1166 | 1136 | 1132 | 1126 | 1229 |
| TrapU | 20573 | 19107 | 4036 | 3888 | 3727 | 3883 | 3890 | 3957 | 4108 |
| TrapW | 21423 | 19766 | 3400 | 3271 | 3162 | NA | NA | NA | 3458 |
| TrapX | 10684 | 9343 | 2857 | 2750 | 2654 | 2553 | 2677 | 2531 | 2907 |
| **Egypt** | | | | | | | | | |
| ALEXAN-1 | 5788 | 5780 | 568 | 561 | 554 | 562 | 564 | 565 | 579 |
| ALEXAN-2 | 8646 | 864 | 625 | 617 | 613 | 613 | 613 | 620 | 627 |
| **Saudi Arabia** | | | | | | | | | |
| SaudiArabia Trap1 | 8691 | 8685 | 678 | 659 | 650 | 659 | 659 | 609 | 691 |
| **Pakistan** | | | | | | | | | |
| Pakistan Museum of Natural History | 17901 | 17900 | 2199 | 2143 | 2088 | 2129 | 2018 | 2112 | 2262 |
| **Honduras** | | | | | | | | | |
| Base Camp | 10409 | 7279 | 1986 | 1935 | 1895 | 1879 | 1876 | 1866 | 1993 |
| Cantiles | 4308 | 3104 | 1094 | 1066 | 1044 | 1060 | 1039 | 1064 | 1104 |
| Cortecito | 1304 | 1169 | 691 | 682 | 674 | 676 | 676 | 671 | 689 |
| Danto | 818 | 602 | 346 | 341 | 339 | 335 | 346 | 359 | 346 |
| GU | 14460 | 12049 | 3509 | 3421 | 3358 | 3367 | 3375 | 3370 | 3540 |

*Includes counts of unsequenced Chironomidae morphospecies (Morphospecies 1: 8595 specimen, Morphospecies 2: 313)

Table 2: Number of species in top 20 most common insect families that are found in single and multiple sites

| **Family** | **# species in one site** | **# species in multiple sites** | **Total** |
| --- | --- | --- | --- |
| Diptera: Cecidomyiidae | 4839 | 48 | 4887 |
| Hymenoptera: Platygastridae | 1466 | 44 | 1510 |
| Diptera: Chironomidae | 952 | 16 | 968 |
| Diptera: Phoridae | 907 | 37 | 944 |
| Hymenoptera: Ichneumonidae | 887 | 16 | 903 |
| Hymenoptera: Braconidae | 842 | 15 | 857 |
| Diptera: Sciaridae | 819 | 26 | 845 |
| Diptera: Ceratopogonidae | 737 | 16 | 753 |
| Hemiptera: Cicadellidae | 610 | 16 | 626 |
| Hymenoptera: Formicidae | 605 | 10 | 615 |
| Hymenoptera: Eulophidae | 564 | 9 | 573 |
| Hymenoptera: Bethylidae | 393 | 5 | 398 |
| Lepidoptera: Erebidae | 371 | 20 | 391 |
| Diptera: Mycetophilidae | 307 | 6 | 313 |
| Hemiptera: Aleyrodidae | 292 | 9 | 301 |
| Diptera: Chloropidae | 269 | 31 | 300 |
| Diptera: Psychodidae | 285 | 12 | 297 |
| Diptera: Muscidae | 262 | 26 | 288 |
| Diptera: Dolichopodidae | 240 | 7 | 247 |
| Diptera: Sphaeroceridae | 191 | 16 | 207 |
| Lepidoptera: Crambidae | 116 | 8 | 124 |

**Table 3:** Robustness of list of top 20 families as found by original analysis vs after merger of families in in top 21-30 with their sister clades. If sister clade was not in the dataset or the group is not monophyletic, taxon was merged all families that together form a monophyletic clade as well as with nearest available clade. Old rank is for taxon in bold. References used are provided below the table.

| **New rank** | **Old rank** | **Taxon** |
| --- | --- | --- |
| 1 | 1 | Diptera: Cecidomyiidae |
| 2 | 2 | Diptera: Ceratopogonidae |
| 3 | 3 | Diptera: Chironomidae |
| 4 | 4 | Hymenoptera: Platygastridae |
| 5 | 5 | Diptera: Psychodidae |
| 6 | 6 | Diptera: Sciaridae |
| 7 | 7 | Diptera: Phoridae |
| 8 | 8 | Hymenoptera: Formicidae |
| 9 | 9 | Hemiptera: Cicadellidae |
| 10 | 10 | Hymenoptera: Ichneumonidae |
| 11 | 11 | Hymenoptera: Braconidae |
| 12 | 12 | Diptera: Dolichopodidae |
| 13 | 13 | Diptera: Muscidae |
| 14 | 30 | Lepidoptera: **Gelechiidae** + Cosmopterigidae^1,2^ |
| 15 | 14 | Lepidoptera: Erebidae |
| 16 | 15 | Diptera: Sphaeroceridae |
| 17 | 16 | Hemiptera: Aleyrodidae |
| 18 | 17 | Hymenoptera: Eulophidae |
| 19 | 18 | Hymenoptera: Bethylidae |
| 20 | 19 | Diptera: Chloropidae |
| 21 | 20 | Lepidoptera: Crambidae |
| 22 | 21 | Coleoptera: **Staphylinidae** + Silphidae + Leiodidae + Agyrtidae^3^ |
| 23 | 23 | Diptera: **Mycetophilidae** + Lygistorrhinidae + Keroplatidae + Bolitophilidae + Ditomyiidae + Diadocidiidae^4**^ |
| 24 | 22 | Diptera: **Limoniidae** + Tipulidae + Cylindrotomidae^5^ |
| 25 | 27 | Coleoptera: **Chrysomelidae** + Cerambycidae + Megalopodidae^3^ |
| 26 | 24 | Coleoptera: **Curculionidae** + Brentidae^3^ |
| 27 | 25 | Psocoptera: **Lepidopsocidae** + Trogiidae + Psoquillidae^6^ |
| 28 | 26 | Diptera: **Drosophilidae** + Cryptochetidae + Braulidae^7^ |
| 29 | 28 | Hymenoptera: **Mymaridae** (sister to rest of Chalcicoidea, including Eulophidae (Rank 17))^8,9^ |
| 30 | 29 | Diptera: **Tachinidae** + Polleniidae^10^ |

****** Alternate hypotheses by Ševčík et al. (2016)^11^ would group Mycetophilidae with a clade containing Sciaridae which is in top 10 families (Rank 6)

**Supplementary Material 1**

Estimation of true global insect species diversity

Stork et al. (2015) use the Ratio between butterfly species diversity and insect diversity in UK based on Barnard (2011). This is extrapolated to 15000-20000 butterfly species present globally. Thus the estimate of 5.4-7.2 million species million is derived from $\frac{67}{24043}\times15,000$ and $\frac{67}{24043}\times20,000$. There are overall 652 Cecidomyiidae described in the UK (2.7% of total diversity). Assuming a 19.98% diversity of Cecidomyiidae, the number of species to remaining to be described can be obtained by solving $\frac{x + 652}{x + 24043}=0.1998$. This estimates that Britain has approx. 5188 unknown Cecidomyiidae species, taking overall diversity to 29231 species. Based on the ratio above, this takes the estimated species diversity to 6.5-8.7 million species.
